# GUNC: Detection of Chimerism and Contamination in Prokaryotic Genomes

**DOI:** 10.1101/2020.12.16.422776

**Authors:** Askarbek Orakov, Anthony Fullam, Luis Pedro Coelho, Supriya Khedkar, Damian Szklarczyk, Daniel R Mende, Thomas SB Schmidt, Peer Bork

**Affiliations:** Structural and Computational Biology Unit, European Molecular Biology Laboratory, 69117 Heidelberg, Germany; Institute of Science and Technology for Brain-Inspired Intelligence, Fudan University, Shanghai, China; Key Laboratory of Computational Neuroscience and Brain-Inspired Intelligence (Fudan University), Ministry of Education, China; Institute of Molecular Life Sciences, University of Zurich, Zurich, Switzerland; Swiss Institute of Bioinformatics, Lausanne, Switzerland; Department of Medical Microbiology, Amsterdam University Medical Center, Amsterdam, The Netherlands; Max Delbrück Centre for Molecular Medicine, Berlin, Germany; Molecular Medicine Partnership Unit, University of Heidelberg and European Molecular Biology Laboratory, Heidelberg, Germany; Department of Bioinformatics, Biocenter, University of Würzburg, Würzburg, Germany

## Abstract

Genomes are critical units in microbiology, yet ascertaining quality in prokaryotic genomes remains a formidable challenge. We present GUNC (the Genome UNClutterer), a tool that accurately detects and quantifies genome chimerism based on the lineage homogeneity of individual contigs using a genome’s full complement of genes. GUNC complements existing approaches by targeting previously underdetected types of contamination: we conservatively estimate that 5.7% of genomes in GenBank, 5.2% in RefSeq, and 15-30% of pre-filtered ‘high quality’ metagenome-assembled genomes in recent studies are undetected chimeras. GUNC provides a fast and robust tool to substantially improve prokaryotic genome quality. Source code (GPLv3+): https://github.com/grp-bork/gunc

## Introduction

Genomes are the genetic blueprint of prokaryotic lineages, a fundamental unit of microbiology [1] at the heart of the ongoing census of the microbial world [2,3] and essential to the study of microbial ecology and evolution [4]. Twenty-five years after the first release of a complete bacterial genome in 1995 [5], more than 700,000 prokaryote genomes have been deposited to NCBI GenBank [accessed 30th of July 2020], doubling almost yearly as genome-based analyses have become the backbone of many disciplines in microbiology. Historically, the vast majority of microbial genomes have been derived from cultured isolates which directly links genome sequences to a physical sample, but excludes the significant number of species that cannot be easily cultivated [4,6].

A promising approach to overcome this deficiency is the delineation of genomes from complex microbial communities using metagenomic data. As early as 2004, nearly complete metagenome-assembled genomes (MAGs) were used to chart the diversity of an acid mine drainage microbial biofilm [7]. Since then, algorithmic advances in binning tools such as canopy clustering [8], CONCOCT [9], MaxBin [10], ABAWACA [11] or metaBAT [12] have enabled the automated recovery of MAGs at large scales, with individual studies now routinely reporting tens of thousands of novel genomes [13–15]. MAGs have led to the discovery of novel deep-branching lineages previously eluding cultivation-based approaches, such as the Asgardarchaeota [16] or the bacterial Candidate Phyla Radiation [11,17], thereby substantially expanding the microbial tree of life [18,19]. Moreover, MAGs can be taxonomically resolved to strain level [20,21] which is particularly beneficial in undersampled environments where reference genomic coverage is scarce [22,23].

As analyses in many microbiological disciplines now critically depend on high-quality genomes, the sheer amount of accruing genomic data, calls for an automated rapid and accurate quality assessment. A substantial fraction of deposited genomes, even those of supposedly high quality in dedicated databases (e.g. Refseq, [24]), contain foreign genome fragments [25] that can originate both *in vitro* and *in silico* (Fig. 1a). Errors in isolate-derived genomes are typically introduced during physical sample processing, *e*.*g*. due to contamination of reagents or culture media [25]. In contrast, the principal error sources in MAGs are expected to be computational [22]: misassembly (i.e., genomic fragments from multiple sources are wrongly assembled together, resulting in chimeric contigs) and mis-binning (contiguous fragments from different sources are erroneously assigned to the same genomic bin, resulting in *chimeric* genomes). Of these two, mis-binning is expected to be the major source of errors, as misassemblies are relatively rare [26]. Genome quality is mainly assessed based on *fragmentation* (*i*.*e*., the size distribution of assembled contigs, with ‘closed’ genomes as the optimum), *completeness* (the fraction of the source genome captured), and *contamination* (‘surplus’ genomic fragments originating from other sources), frequently estimated based on ubiquitous and single-copy marker genes (SCGs), using tools such as BUSCO [27] or CheckM [28].

**Fig. 1.**
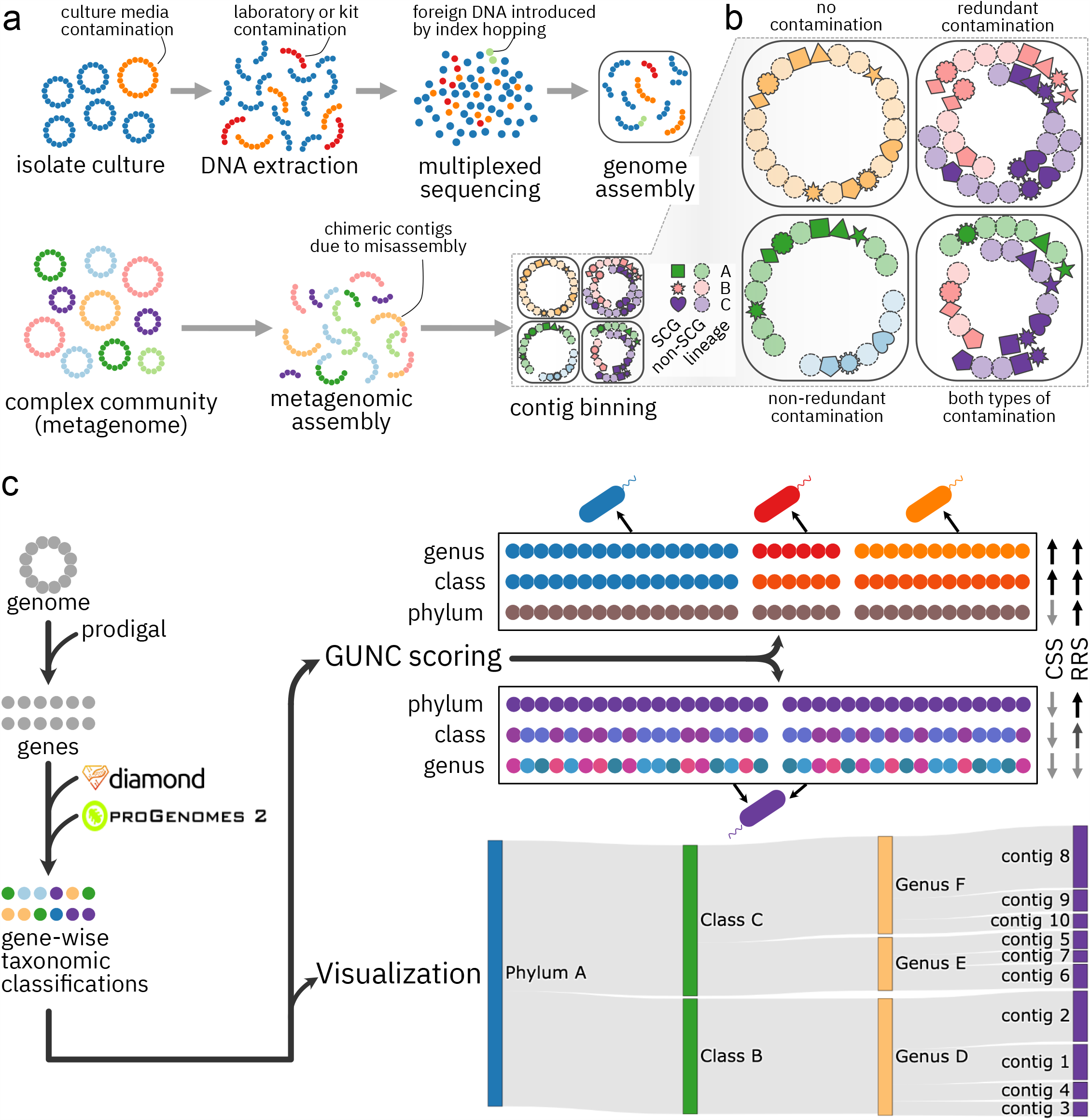
GUNC quantifies chimerism in prokaryotic genomes. **a** Genome contamination may originate *in vitro* (e.g., from culture media, laboratory equipment or kits, index hopping during multiplexed sequencing) or *in silico* (contig misassembly, erroneous binning). Genomes are represented as circular chromosomes, contigs as sequences of genes (dots). **b** Two types of genome contamination can be distinguished operationally: *redundant* contamination by surplus genomic material (‘more of the same’) and *non-redundant* contamination by non-overlapping fragments from unrelated lineages (‘something new’, e.g. novel sets of orthologues). Different single-copy marker genes (SCGs) are shown as solid shapes, other genes as dashed circles; colours indicate different source lineages. **c** GUNC workflow. For a given query genome, genes are called using prodigal, then mapped to the GUNC reference database (based on proGenomes 2.1) using diamond to compute GUNC scores and to generate interactive Sankey diagrams to visualize genome taxonomic composition. GUNC quantifies genome chimerism and reference representation across taxonomic levels. Clade separation scores (CSS) are high if gene classification to distinct lineages (represented by different colours) follows contig boundaries. Reference representation scores (RRS) are high if genes map closely and consistently into the GUNC reference space. The top example illustrates a chimeric genome with good reference representation; the bottom example a non-contaminated genome that is not well represented in the GUNC reference.

Erroneous genomes can affect analyses in different ways: whereas type II errors introduced due to missing or truncated genetic elements in fragmented or incomplete genomes can usually be mitigated, contaminating fragments are detrimental to biological interpretation, as they may cause false inferences about a genome’s functional repertoire or structure [22,25]. Operationally, two types of genome contamination can be distinguished (Fig. 1b). *Redundant* contamination involves surplus genomic fragments (e.g., genes from gene families already present in the focal genome) from related sources (often within the same or related lineages). In contrast, *non-redundant* contamination involves foreign fragments (e.g., gene families not encoded in the focal genome’s lineage) that replace or extend part of the source genome with unrelated, non-overlapping material, leading to chimeric genomes. Intuitively, redundant contamination can be thought of as an addition of ‘more of the same’ genomic material, whereas non-redundant contamination adds ‘something new’.

Genome contamination can be difficult to estimate, particularly for phylogenetically novel lineages that are not well represented by existing references. By design, SCG-based estimators of genome quality can detect redundant contamination with high sensitivity, but they are less sensitive towards non-redundant contamination, since they only consider inventories of expected SCGs as a whole, remaining agnostic to conflicting lineage assignments between individual genes [28,29]. The widely used CheckM algorithm [28] first places a query genome into a reference phylogeny, then defines a clade-specific set of expected SCGs to estimate genome completeness and contamination. However, for genomic chimeras of unrelated lineages, phylogenetic placement will be more conservative, nearer the root, limiting the range of consensus SCGs: in the extreme case of root-level placement, quality estimates are based on only 43 near-universal genes [28], corresponding to just 1-2% of an average prokaryotic genome. Moreover, lineage-specific SCGs, in particular those with deep phylogenetic roots, are often not evenly distributed across the genome, but locally clustered [30,31], additionally limiting their representation of the query genome. Such biased quality estimates can have a detrimental impact on biological interpretation, as demonstrated in the case of the novel deeply branching lineage Rokubacteria [29], and more recently shown anecdotally for manually curated human gut-derived MAGs [22]. As a result, the use of SCG inventories as the exclusive estimator of genome quality has been questioned, in particular in the context of large-scale automated genome binning [22].

Here we present GUNC (the Genome UNClutterer), a fully integrated workflow to estimate genome chimerism based on the full gene complement, using an entropy-based measure of lineage *homogeneity* across contigs. In various simulation scenarios, we demonstrate that GUNC accurately quantifies genome contamination at high sensitivity and can pinpoint problematic contigs within genome bins. We further identify a substantial fraction of chimeric genomes in GenBank [32], the Genome Taxonomy Database [33] and recently published large-scale MAG datasets [13–15] for which contig misbinning also leads to inflated estimates of phylogenetic diversity and taxonomic novelty. GUNC source code is available under a GPLv3 license at https://github.com/grp-bork/gunc.

## Results

### GUNC estimates genome quality based on contig homogeneity using the full complement of genes

Our rationale in designing GUNC was to estimate genome quality based on the phylogenetic homogeneity of contigs with respect to a genome’s entire gene content. Using a filtered and curated set of high quality reference genomes derived from proGenomes2.1 [34], GUNC infers each gene’s clade membership across a hierarchy of taxonomic levels, using taxonomy as a proxy for phylogeny. While ideally a genome would receive a single dominant label, corresponding to its true classification, there are two main reasons why this may not happen: it may be contaminated or it may lie outside the reference set and thus, due to prediction limitations, receive a set of inconsistent inferences.

GUNC attempts to distinguish between these two cases by computing a *clade separation score* (CSS) which builds upon an entropy-based metric [35]. The CSS measures how diverse the taxonomic assignments are within each contig, normalized to the diversity across the whole genome and to the expected entropy when there is no relationship between taxonomy and contig labels, thus returning a value between 0 and 1 (see Methods). Intuitively, if a genome is composed of contigs that are internally homogeneous, but disagree with each other, then the metric will return a value closer to 1. On the other hand, a genome that, because it lies outside the reference set, is assigned a myriad of labels, but where the labeling does not follow contig boundaries, will have a CSS closer to 0. The CSS quantifies the degree to which a genome is a *chimeric* mixture of distinct lineages following non-random distributions across contigs. GUNC computes CSS values at all major taxonomic levels and can thus indicate the approximate phylogenetic depth at which distinct source genomes diverged.

Importantly, the CSS is a measure of confidence when labelling a genome as chimeric, and is sensitive even to small portions of contaminant if these are well circumscribed by contig boundaries. GUNC additionally quantifies the fraction of genome contamination in two ways: as the fraction of total genes assigned to non-major clade labels (GUNC contamination), and as the “effective number of distinct clades” in a genome, based on the Inverse Simpson Index of the clade size distribution (see Methods). To assess a genome’s quality based on GUNC, both GUNC contamination and CSS should be taken into account.

Finally, GUNC also estimates how closely a query genome is represented by the underlying reference set. The *reference-representation score* (RRS, see Methods) is based on the average identity of query genes to the reference and the number of spurious mappings, in order to further inform the interpretation of the CSS and genome contamination. Beyond mere statistics, GUNC also provides interactive visualisations of a query genome’s taxonomic composition as alluvial Sankey diagrams at gene-level resolution (see Methods). A full overview of the GUNC workflow is provided in Fig. 1c.

### GUNC accurately quantifies genome contamination in multiple simulated scenarios

We benchmarked and validated GUNC in multiple scenarios, simulating various degrees of genome chimerism, source genome relatedness and reference representation (see Fig. 2a and Methods). All simulated genomes were generated from a curated high-quality set derived from proGenomes2.1 [34], which is also the basis for the default reference set used by GUNC (Fig. 2a ‘type 1 genomes - in reference’). We simulated decreasing reference representation by iteratively removing entire clades from the GUNC training set at varying taxonomic levels (‘type 2 genomes - out of reference’). Type 1 genomes were used as the contamination-free baseline in the subsequent benchmarks.

**Fig. 2.**
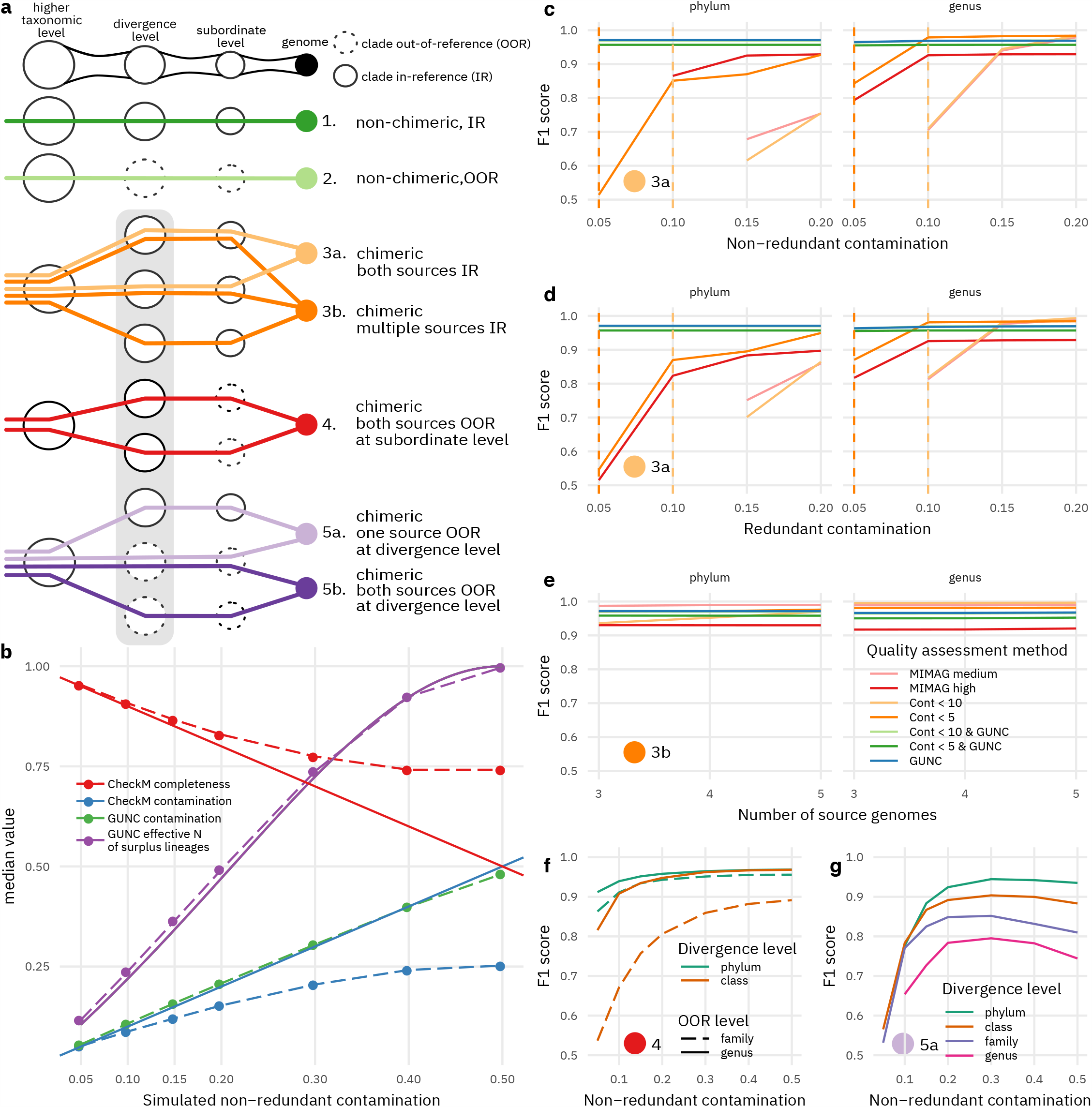
GUNC accurately detects chimerism in incrementally challenging simulation scenarios. **a** Overview of different types of simulation scenarios. Genomes (filled circles) were simulated as mixtures of lineages (horizontal lines) diverging at various taxonomic levels (columns) from clades (void circles) contained in the GUNC reference (solid lines) or not (dashed). See Methods for details. **b** Median CheckM completeness and contamination estimates (dashed lines) diverged from true values (solid lines) with increasing levels of simulated contamination (type 3a in panel a), whereas GUNC estimates of contamination (green; theoretically expected values as blue solid line) and effective number of surplus lineages (purple) were highly accurate. See Fig S1 for an equivalent plot on type 3b genomes. **c-f** Detection accuracy across simulation scenarios, quantified using F1-scores (y axis) across increasing levels of simulated contamination (x axis). Data shown for scenarios 3a (c-d), 3b (e), 4 (f) and 5a (g); full panels for types 3a and 3b in Fig S2. MIMAG criteria were defined as CheckM contamination <10%, completeness >50% (medium) and contamination <5%, completeness >90% (high); note that MIMAG criteria on rRNA and tRNA presence were not applied; Cont, CheckM contamination; GUNC, default GUNC CSS cutoff of >0.45.

We simulated chimeric genomes by mixing fragments from two (‘type 3a’) or multiple (‘type 3b’) sources, varying the taxonomic level at which sources diverged (‘divergence level’), but retaining source lineage representatives in the reference training set (see Methods). Non-redundant contamination was simulated by replacing part of an acceptor genome by a size-matched contaminant fraction of a donor genome; to simulate redundant contamination, surplus donor fragments were added to complete acceptor genomes. We observed that CheckM systematically overestimated completeness and underestimated contamination for genomes with simulated non-redundant contamination (Fig. 2b), largely independent of the taxonomic level of source genome divergence. This bias further increased when two source genomes were mixed at more equal shares (Fig. 2b) and followed a similar trend when multiple source genomes were mixed (Fig. S1). In contrast, GUNC accurately estimated the contamination and the effective number of surplus lineages represented in the query genome.

This difference in quantitative estimates translated into differential accuracy in a binary classification of genome quality, assessed using the harmonic mean of precision and recall (F1 score) at different levels of contamination (Fig. 2c-e). GUNC scores at default thresholds (CSS <0.45, contamination filtered at <2%; see Methods and Figure S2) generally outperformed the widely used MIMAG standard parameters [36] for medium (contamination < 10%, completeness ≥ 50%) and high (contamination < 5%, completeness > 90%) genome quality when based on CheckM estimates. GUNC accurately detected both non-redundant (Fig. 2c) and redundant contamination (Fig. 2d) for mixtures of two or more (Fig. 2e) source genomes (Fig. 2e), with F1 scores of ≥0.96 even at simulated contaminant fractions of just 5%. As expected, GUNC accuracy was consistently high with the only exception of species-level chimerism where it performed suboptimally at lower portions of contamination. In contrast, CheckM-based classification was less accurate for phylogenetically deep chimeras, dropping as low as F1 < 0.5 (F-scores below 0.5 not plotted). Interestingly, including the completeness criterion (in MIMAG medium and high thresholds) provided only mild performance improvements in our simulations when compared to classification based only on CheckM contamination. A strict CheckM contamination threshold of <5% slightly outperformed GUNC for species-level chimeras (Fig. S2a-c), while also occasionally showing minute performance benefits at much higher degrees of contamination (≥20%) for higher taxonomic levels, as GUNC performance generally plateaued at 100% sensitivity with a low fraction of residual false positive calls.

The simulation scenarios of types 3a & 3b (Fig. 2c-e) assume that the lineages, but not the genomes themselves, of both contaminant and acceptor are represented in the reference training set. In practice, however, this is often not the case, in particular for novel lineages and MAGs. We therefore simulated scenarios of genome chimerism between source lineages that are themselves out of reference (types 4, 5a & 5b in Fig. 2a). Note that by design, such leave-one-out simulations are not possible with CheckM, as the pre-curated reference phylogeny and marker gene sets included with the software cannot be modified accordingly. Genomes of type

4 simulated chimerism between deeply branching source lineages with limited reference representation at the divergence level and none at subordinate levels. For example, these genomes represent chimeras of two novel families or genera within distinct previously described phyla or classes. GUNC accurately detected such chimeras, even at low fractions of contamination (5-10%; Fig. 2f). We next simulated even more challenging scenarios in which one (‘type 5a’) or both (‘type 5b’) source lineages were not represented even at the level of divergence, corresponding to chimeras of entirely novel lineages. GUNC accurately detected ‘type 5a’ at contaminations ≥10% at phylum to family level (Fig. 2g), though performance deteriorated towards lower contamination portions and shallower phylogenetic novelty (Fig. S2).

As expected, GUNC was not able to accurately detect chimerism in scenario ‘5b’, i.e. if the clades of both source lineages were out of reference. Instead, GUNC addresses the challenges posed by novel lineages, both as possible contaminants and as units of discovery, via reference representation scores (RRS) across taxonomic levels, based on the average identity of query genes to their closest reference counterparts (see Methods). High RRS values indicate that genomes map confidently into the reference space at a given taxonomic level, whereas low RRS indicate phylogenetic novelty. Using simulations, we confirmed that GUNC RRS can predict the taxonomic level at which a query genome is novel (Fig. S3), in particular for deeply branching novel lineages (F1=0.98 for novel phyla), and regardless of whether the query is itself chimeric (type 5b) or not (type 2). We suggest that the CSS and RRS be used in conjunction to assess genome quality, depending on the expected phylogenetic novelty in the dataset under investigation.

### Extensive undetected contamination among reference genomes and MAGs

We calculated GUNC scores for various public datasets of both isolate-derived and metagenome-assembled genomes to detect hitherto overlooked genome chimerism (Fig. 3a, see Fig. S5 for alternative GUNC parameters). We applied default GUNC thresholds (CSS > 0.45, see Methods), conservatively ignoring species-level chimerism, i.e. only considering chimerism between genomes involving distinct genera and higher taxonomic ranks. The resulting GUNC profiles for 1,375,848 genomes are available as Supplementary Data (see availability of data and materials section). Using these parameters, 5.7% of 701,698 prokaryotic genomes in GenBank [accessed 30th of July 2020] [32] and 5.2% in the more restrictive RefSeq [24] were flagged as potentially chimeric. Genomes annotated as ‘environmental’ or ‘metagenome-derived’ (i.e., MAGs) were substantially enriched for chimeras in GenBank, accounting for 18.4% of chimeric genomes even though the overall GenBank MAG share was only 9.4%. Moreover, by following up genome taxonomic annotations, we observed that GenBank contains ‘cryptic’ MAGs that were not annotated as metagenome-derived by submitters. Indeed, for proGenomes 2.1 [34], a more vetted and curated GenBank subset totalling 84,095 genomes, the fraction of flagged genomes was only 3.6%.

**Fig. 3.**
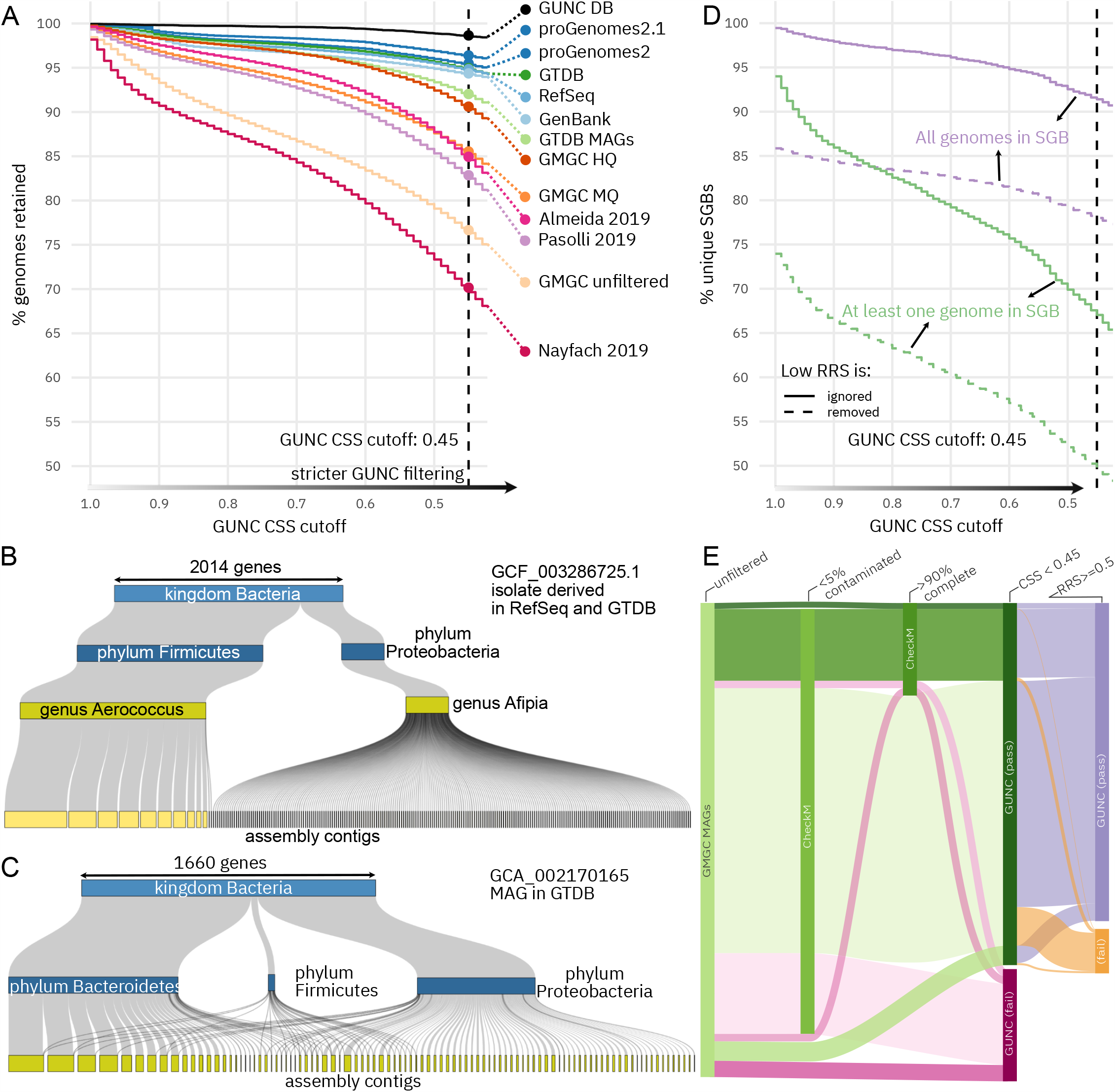
Extensive undetected chimerism in public genome databases and large-scale MAG datasets. **a** Cumulative plots summarizing genome quality for various genome reference and MAG datasets. The y axis shows the fraction of genomes passing GUNC filtering at increasing stringency (x axis), up to the default CSS threshold of 0.45, conservatively ignoring species-level scores. Note that the Almeida, Pasolli and Nayfach sets were pre-filtered using variations of the MIMAG medium criterion based on CheckM estimates. GTDB, Genome Taxonomy Database; GMGC, Global Microbial Gene Catalogue. **b** Example of detected contamination in an isolate-derived reference genome for which around one fifth of genes were assigned to a different phylum, scattered across hundreds of small contigs. **c** Example of detected contamination in a MAG for which genes assigned to two major different phyla were well separated into distinct contigs. **d** Cumulative plots summarizing the quality of species-level genome bins (SGBs) defined by Pasolli et al. 2019. Lines indicate the fraction of SGBs (y-axis) containing at least one or exclusively chimeric genomes at increasingly stringent GUNC cutoffs (x axis) conservatively ignoring species-level scores. For both series, intervals correspond to edge scenarios in which genomes with limited reference representation are either conservatively ignored (treated as non-chimeric, upper lines) or aggressively removed (lower lines); the true fraction of chimeric SGBs likely falls in between. **e** Differential filtering of MAGs in the GMGC set based on CheckM contamination (<5%), CheckM completeness (>90%) and GUNC (CSS <0.45, ignoring species-level scores).

Among the flagged isolate genomes in GenBank, RefSeq, proGenomes 2.1 and the Genome Taxonomy Database (GTDB, [2]), we frequently observed patterns consistent with biological contamination, e.g. of culture media or reagents. For example, Fig. 3b shows an isolate of the Firmicute *Aerococcus urinae*, contaminated by a *Afipia broomeae* (phylum Proteobacteria) scattered across many small contigs. The division between both source genomes clearly follows contig boundaries, indicating that the highly fragmented *Afipia* genome may have been partially assembled from a lowly abundant contamination co-sequenced at low coverage (Table S1).

While contamination in isolate genomes was usually restricted to small contigs due to the low abundance of the contaminant species, the sizes of contaminant contigs in chimeric MAGs were more evenly distributed, (cf Fig. 3c for a phylum-level chimera MAG from a marine metagenome [41]).

As observed in GenBank, genome chimerism was more common among MAGs than among isolate genomes. The GTDB comprises both types, extending a GenBank-derived core set with automatically generated MAGs for underrepresented lineages, all filtered based on CheckM quality estimates and clustered into species-level units by average nucleotide identity. Among the GTDB, MAGs were more prone to being contaminated (8.0%) than single-cell derived (6.7%) or isolate genomes (4.6%). Chimeric genomes likely inflate estimates of total phylogenetic diversity in the GTDB: 1,009 (3.2%) of species-level clusters in the GTDB consisted entirely of contaminated genomes, and a further 1,760 (5.6%) contained at least one.

For the human gut, three recent studies alone generated hundreds of thousands of MAGs by assembling and locally binning metagenomic data from several thousand samples [13–15]. All three teams relied on methodologically largely equivalent approaches for MAG generation and filtering, using variations of the MIMAG ‘medium’ quality standards, based on CheckM estimates. GUNC identified 17.2%, 15.1% and 29.9% of the pre-filtered Pasolli, Almeida and Nayfach MAG sets as putatively chimeric at genus level or above (Fig 3a), revealing extensive levels of previously undetected non-redundant contamination. Among the species-level genome bins (SGBs, clustered at 95% average nucleotide identity) described by Pasolli et al [13] chimeric genome and 8.5% consisted entirely of chimeras (Fig. 3d), with even higher rates among ‘novel’ SGBs (not containing any reference genomes) and small clusters: 18% of ‘novel’ singleton SGBs were formed by chimeric MAGs. Thus, the 17.7% of chimeric genomes in the Pasolli set may have strongly impacted both SGB clustering and, as a consequence, biological interpretation.

To further quantify the differential effects of CheckM and GUNC filters on MAG datasets, we re-analysed the 278,629 MAGs derived from the Global Microbial Gene Catalog dataset (Coelho et al, *in revision*). GUNC flagged 23.4%, 14.5% and 9.4% of the raw, MIMAG ‘medium’ and MIMAG ‘high’ quality filtered GMGC MAGs as chimeric, respectively, comparable to levels in other tested pre-filtered MAG sets (Fig. 3a). The CheckM 5% contamination criterion was highly permissive, flagging just 10.3% of all GMGC MAGs. GUNC was more restrictive, flagging 23.4% of total genomes and 20.5% of genomes passing the CheckM contamination filter (reciprocally, only 7.0% of genomes passing the GUNC CSS filter were flagged at CheckM contamination >5%). Overall, CheckM and GUNC contamination filters agreed on 76% of genomes, at a Pearson correlation between nominal contamination estimates of 0.2. The CheckM completeness criterion, capturing an entirely orthogonal signal, was the overall most restrictive filter, flagging 80% of genomes as ≤90% and 50% as ≤50% complete (Fig. 3e and S6). Relaxing the completeness criterion further pronounced the differential impact of GUNC and CheckM contamination filters (Fig. S6), with GUNC being consistently more sensitive.

## Discussion

Chimerism and contamination can have considerable impact on the biological interpretation of a genome, in particular by causing false inferences about phylogenetic placement and functional repertoires [22,25], thus there is a need for fast and accurate methods for automated genome quality control. As we have shown, GUNC quantifies even small levels of chimerism in prokaryotic genomes, with robust performance even if one or multiple source lineages of a composite genome are not well represented in the GUNC reference database.

GUNC is designed to complement existing estimators of prokaryote genome quality such as the *de facto* standard in the field, CheckM [28], and addresses error types that elude marker gene-based methods. GUNC represents a genome as its full gene complement, not just as an inventory of ‘expected’ core genes and is therefore robust to common artefacts resulting from erroneously conservative phylogenetic placement. Moreover, the enhanced resolution of a gene-centric genome representation has been shown to increase accuracy for related problems, such as e.g. taxonomic classification [42,43]. GUNC can provide gene-level resolution even for composite genomes of deeply branching source lineages, a type of chimeras that are notoriously difficult to detect automatically as sets of shared marker genes rapidly shrink with increasing phylogenetic depth. We demonstrated that GUNC scoring was highly accurate in incrementally challenging simulation scenarios. Moreover, GUNC quantifies the ‘novelty’ of a query genome relative to its reference set, thus further qualifying quality estimates, as confidence decreases along with reference representation. Nevertheless, as demonstrated in incrementally challenging simulation scenarios, GUNC accurately detects chimerism even among novel lineages.

By design, GUNC does not quantify genome completeness, as it does not attempt to infer an expected set of a lineage’s core genes. This also means that GUNC does not attempt to resolve redundant contamination between very closely related (or even identical) lineages – a use case at which marker gene-based methods excel. By the same token, we caution against an over-interpretation of GUNC CSS and contamination estimates at species or strain resolution: GUNC’s underlying gene-wise taxonomy assignments become less precise between closely related lineages that share substantial genetic material at very high sequence similarity, potentially causing an overestimation of contamination. Moreover, very closely related lineages are prone to recombination and exchange of genomic material which can further confuse gene level classifications. Nevertheless, GUNC reports scores at all taxonomic levels and in practice accurately detects species-level chimerism (see Figure S2 and Figure S8).

Applying permissive GUNC default thresholds, we demonstrated that a substantial fraction of genomes in public repositories show clear contamination signatures that were not picked up previously. As expected, metagenome-assembled genomes were much more prone to chimerism than those derived from isolates, irrespective of a lineage’s novelty relative to the GUNC reference. Among four large-scale datasets of automatically generated MAGs from human microbiomes [13–15] and various environments (Coelho et al, *in revision*), we found extensive levels of undetected genome contamination, with a disproportionate impact on estimates of phylogenetic novelty.

As GUNC complements existing tools to estimate genome quality, using orthogonal information to address types of genome contamination that are currently overlooked, a combination of filters based on genome fragmentation, CheckM completeness, CheckM contamination and GUNC may greatly refine automatically generated MAG datasets. GUNC offers a dedicated workflow to accomplish this, integrating CheckM results with GUNC scores for nuanced estimates of genome quality at high throughput. We expect that an automated, rapid and accurate quantification of genome contamination will further enable genome-centric microbiology at large scale and high resolution.

## Methods

### GUNC workflow and implementation

The core workflow of GUNC consists of three modules (see Figure 1C). First, for any query genome, genes are called using *prodigal* [44], although per-gene protein sequences can alternatively be supplied by the user directly. Protein sequences are then mapped against representative genomes in the GUNC database (derived from species-representative genomes in proGenomes 2.1 [34]) using *diamond* [45], retaining best hits (-k 1) without applying an evalue filter (-e 1) as alternative filtering is applied downstream. Annotated plasmids and other non-chromosomal genomic elements are excluded from the reference to reduce nonspecific hits between lineages within plasmid host range. Moreover, the reference set was semi-manually curated, removing clear cases of genomic chimerism.

For each query gene, taxonomic annotations at 7 levels (kingdom, phylum, class, order, family, genus, species) are inherited from the best hit via the manually curated proGenomes 2.1 taxonomy. To filter against mapping noise, taxonomic clade labels recruiting less than 2% of all mapped genes are dropped. GUNC scores (see below) are then calculated based on inferred taxonomic labels, query gene contig membership, sequence identity to database hits and the fraction of mapped and filtered hits. Finally, GUNC offers a visualization module to automatically generate interactive Sankey alluvial diagrams of contig-level taxonomic annotations to enable manual curation and exploration of flagged genomes.

GUNC is implemented in Python3, all code is open source and available at https://github.com/grp-bork/gunc and through bioconda [46] under a GPLv3+ licence. Based on database size and resource requirements, GUNC can be run locally on a personal computer but is also highly parallelizable in a cluster environment.

### Calculation of GUNC scores

GUNC computes several scores to quantify a query genome’s quality, its representation in the GUNC reference database and its levels of putative contamination. The GUNC *clade separation score* (CSS) is an entropy-based clustering measure to assess how homogeneously taxonomic clade labels (T) are distributed across a genome’s contigs (C). It is inspired by the uncertainty coefficient [35],

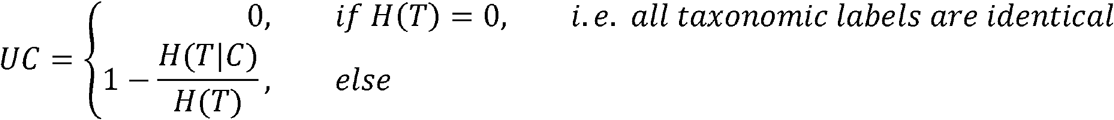

A simple estimator for this quantity is the plugin estimator where C is a set of contigs, T is a set of taxonomic clades, *a*_*ct*_ is a number of genes located in contig *c* and assigned to taxonomic clade *t*, N is the total number of genes in a genome.

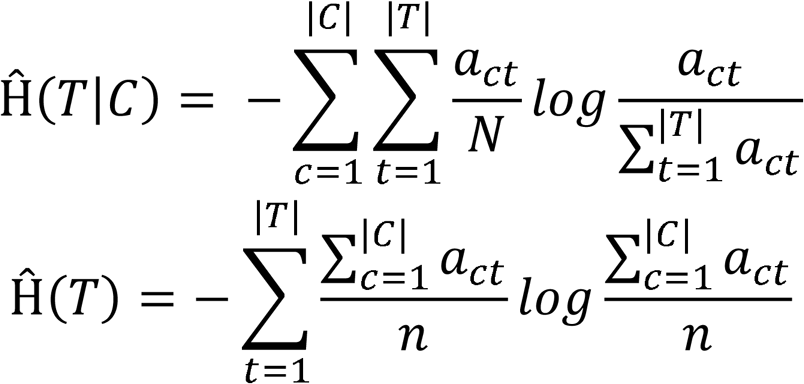

However, this estimator is known to be biased when the number of samples is small [47] and adjusting it for chance leads to more interpretable quantities [48]. In our case, the sums range over the genes in each contig, and, in fragmented genomes, many contigs can contain only a small number of genes. Therefore, we normalize the estimated conditional entropy by the expected value of this estimation under a null model, leading to CSS = 1 - Ĥ(T|C)/Ĥ(T|R),

where Ĥ(T|R) is the expected value of Ĥ(T|C) keeping the same contig size distribution and assuming no relationship between contig membership and taxonomic assignment (in the special case where Ĥ(T|C) > Ĥ(T|R), we set CSS to zero).

The CSS is 0 if the frequency distribution of taxonomic labels in every individual contig exactly follows that across the entire genome. It is 1 if all contigs are ‘taxonomically pure’, i.e. if the distribution of taxonomic labels follows contig boundaries. GUNC outputs CSS scores for every tested taxonomic level, so that users can infer the approximate phylogenetic depth at which source lineages diverged. By default, GUNC adjusts CSS to 0 at every level separately when the portion of called genes left after removal of minor clades, i.e. genes retained index < 0.4, because in that case there are too few remaining genes to calculate scores on at that level. Then, GUNC flags a genome as putatively contaminated if the “adjusted” CSS > 0.45 at any taxonomic level, a threshold benchmarked in a series of simulation scenarios (Figure S7).

The CSS does not carry information about the scale of contamination (i.e., the fraction of contaminant genome), but about the confidence with which a query genome may be considered chimeric. In other words, the CSS assesses whether a genome is contaminated or not, but not how large the contaminant fraction is. GUNC instead quantifies the scale of contamination at each tested taxonomic level using two measures. The total fraction of genes with minority clade labels after filtering (‘GUNC contamination’) is an estimate of the total fraction of contamination in the query genome. Note that this definition differs from that commonly used by tools such as CheckM: designed to quantify non-redundant contamination, GUNC scales by the total query genome size, whereas CheckM estimates redundant contamination by scaling against a theoretical ‘clean’ source genome with a single set of SCGs. In practice, this means that GUNC contamination never exceeds 100%, whereas

CheckM contamination estimates the number of (complete) surplus genomes. GUNC further provides a combined estimate of redundant and non-redundant contamination as the *effective number of surplus clades* (T_eff_) in a query genome, calculated as the Inverse Simpson Index minus 1 (as 1 genome is expected):

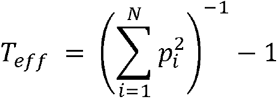

where p_i_ is the fraction of genes assigned to clade i. T_eff_ scales in [0, ∞] and can be interpreted as the number of surplus clades in the query genome considering the weighted contributions of all source lineages.

Finally, GUNC computes a *reference representation score* (RRS) based on the total fraction of genes mapping to the GUNC database (*Portoin*_*genes mapped*_), the fraction of genes retained after noise filtering (*Portoin*_*genes retained*_) and their average similarity to the reference (*Identity*_*mean*_):

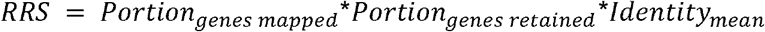

The RRS captures the expectation that out-of-reference genomes will map to the reference to a lower degree (*Portoin*_*genes mapped*_) and at lower similarity (*Identity*_*mean*_). Moreover, among simulated out-of-reference genomes, we empirically observed a characteristic pattern of noisy, low confidence hits scattered unspecifically across multiple clades at very low frequencies; in the RRS, this signature is formalised as the term Portion_genes retained_. High RRS values indicate that a query genome maps well within the GUNC reference space, whereas low RRS indicates poor reference representation to qualify the interpretation of CSS and contamination estimates.

In general, the lower the RRS, the higher the risk of type 1 errors based on CSS (falsely labelling genomes as contaminated): this way, GUNC asserts that genome quality is only confidently estimated where sufficient data is available, and that genomes potentially representing deeply branching novel lineages beyond the GUNC reference are flagged for further (manual) inspection.

### Construction of artificial genomes under different scenarios

Artificial genomes were constructed to simulate different scenarios of genome contamination and reference representation (see Fig. 2a). All simulations were performed using genomes in the curated and taxonomically annotated proGenomes 2.1 database [34], serving as a baseline for clean, in-reference genomes (‘type 1’ in Fig. 2a). Further simulation scenarios are described below. Unless otherwise indicated, simulations were conducted separately for each taxonomic level and at contamination portions of 5%, 10%, 15%, 20%, 30%, 40% and 50%, with 3,000 iterations/genomes per each taxonomic level and contamination portion. In each simulated genome, source genome contigs were randomly fragmented such that contig size was inversely proportional to contig frequency, parameterized based on the empirical frequency-size distributions of MAGs in the Pasolli, Almeida and Nayfach datasets [13–15]. Simulated genomes were then generated from these simulated contigs based on the rules set out below:

**Type 1**. Clean (non-contaminated) genomes, in reference. Taken from progenomes2.1.

**Type 2**. Clean (non-contaminated) genomes, out of reference. Simulated by removing a genome’s entire source lineage from the reference.

**Type 3a**. Binary chimeric genome from two sources, both in reference. Simulated by randomly selecting ‘donor’ and ‘acceptor’ genomes whose lineages diverged at any of the seven tested taxonomic levels (*divergence levels*). A fraction of the acceptor genome was either *replaced* by a matching fraction of donor genome (to simulate non-redundant contamination), or the corresponding fraction of donor genome was *added* to the complete recipient genome (to simulate redundant contamination).

**Type 3b**. Chimera of multiple (3, 4, or 5) source genomes, all in reference. Source genomes from different source clades were mixed at equal shares totaling 1 altogether, e.g. ⅓, ¼ or ⍰ each.

**Type 4**. Binary chimera, both source lineages out of reference at subordinate levels. Source lineage clades removed at subordinate levels (e.g., genus or family) but sister clades retained in reference within the same parent clades (e.g., class or phylum), so that both higher-level source clades were represented at divergence level. Simulated 10,000 times for each taxonomic and contamination level.

**Type 5a**. Binary chimera, one source lineage in reference, one out of reference at divergence level. Recipient genome (in reference) partially replaced by donor genome (out of reference at divergence level).

**Type 5b**. Binary chimera, both source lineages out of reference at divergence level, e.g. no genome available from entire clades (at divergence level) containing source genomes.

### Reanalysis of public datasets

Pasolli et al. [13]: genome fasta files were accessed from http://segatalab.cibio.unitn.it/data/Pasolli_et_al.html on 2020-03-12. SGB annotations of genomes were taken from article supplementary files.

Almeida et.al. [15]: genome fasta files were accessed from ftp://ftp.ebi.ac.uk/pub/databases/metagenomics/umgs_analyses/ on 2020-03-12.

Nayfach et.al. [14]: genome fasta files were accessed from http://bit.ly/HGM_all_60664_fna-OHGM_v1.0_all_60664_fna.tar.bz2 on 2020-03-12.

GenBank genomes were accessed on 30.07.2020. GUNC results for 699,994 GenBank genomes were produced. RefSeq genome set was subsetted from GenBank. Genomes annotated as “derived from metagenomes” or “derived from environmental source” in the “excluded from refseq” column of the GenBank assembly metadata were considered as MAGs.

GTDB [2] genome metadata was accessed from https://data.ace.uq.edu.au/public/gtdb/data/releases/release95/95.0/ on 2020-08-17. Genomes were mapped to GenBank via GenBank accession IDs. Genomes annotated as “derived from metagenomes” or “derived from environmental sample” in the “ncbi_genome_category” column of the metadata table were considered as the GTDB MAGs subset.

The unfiltered set of 278,629 MAGs of the Global Microbial Gene Catalog dataset (GMGC, Coelho et al, *in revision*) is available at gmgc.embl.de.

## Availability of data and materials

The datasets generated and/or analysed during the current study are available in the GUNC webpage, gunc.embl.de/datasets

## Funding

This work was partially supported by EMBL, the German Federal Ministry of Education and Research (grant numbers: 031L0181A; 01Kl1706), the H2020 European Research Council (ERC-AdG-669830) and the German Network for Bioinformatics Infrastructure (de.NBI #031A537B).

### Acknowledgements

The authors thank Chris Creevy for insights on phylogenetically deep horizontal gene transfer. We thank Oleksandr M Maistrenko for comments and feedback on the manuscript, as well as all members of the Bork lab for insightful discussions.

## Author Contributions

A.O. & T.S.B.S. conceived the study and designed analyses. L.P.C., D.S., D.R.M. & P.B. contributed to formulating and implementing core concepts in GUNC scoring. S.K. & D.R.M. provided, curated and analysed reference and test datasets. A.O. performed simulations, benchmarks and analyses, with support by A.F.. A.O., T.S.B.S & P.B. analysed the data, with input from all authors. A.F implemented and optimized the tool for distribution. T.S.B.S. & A.O. wrote the manuscript, with input from all authors. All authors read and approved the final manuscript.

## Figure Captions

***Fig. S1***.

Comparison of median scores from GUNC and CheckM of simulations of genomes type 3a and 3b where source genomes make equal contributions summing 1 in total (e.g. 0.2 from each of 5 sources or 0.25 from each of 4 sources). This shows that the trend from Fig. 2b persists when multiple source genomes are mixed in a simulated chimeric genome.

***Fig. S2***.

F-scores of distinction between clean and chimeric genomes across all divergence levels of source genomes for different simulation scenarios. MIMAG medium is CheckM contamination < 10% and CheckM completeness >50%. MIMAG high is CheckM contamination <5% and CheckM completeness >90% and due to irrelevance to our simulations we decided that additional criteria of presence of rRNAs and tRNAs can be ignored here. “Cont” stands for CheckM contamination and GUNC means GUNC CSS of <0.45 or GUNC contamination <2%. **a** type 3a non-redundant contamination. **b** type 3b redundant contamination. **c** type 3b.

***Fig. S3***.

**a** ROC-curves and AUCs of separation of genomes in-reference (type 1) from genomes out-of-reference (types 2 and 5b) at different out-of-reference levels (faceting) using GUNC reference representation scores (RRS) at matching taxonomic levels. **b** F1-scores (y-axis) of separation of types 2 & 5a from type1 across different RRS cutoffs (x-axis) different out-of-reference (OOR) levels (colors) at which these genomes have no reference representation. GUNC scores at the taxonomic level identical to OOR level were used. Vertical lines indicate RRS scores with highest F1-scores at each OOR level. Cutoff of RRS <0.5 is used to label genomes as “OOR” irrespective of the taxonomic level of max CSS.

***Fig. S5***.

**a** Cumulative plot summarizing genome qualities of various sets of genomes represented by lines of different colors. Any point in a plot indicates a portion of genomes retained in a set (y-axis) after filtering out genomes with GUNC CSS higher than the cutoff (x-axis) & GUNC contamination >5% (ignoring species level scores). **b** Cumulative plot illustrating the number of species-level genome bins (SGBs) (from Pasolli et al. 2019). Lines indicate the portion of unique SGBs retained (y-axis) after filtering out SGBs where either “all” or “at least one” genome has GUNC CSS score higher than the cutoff (x-axis) & GUNC contamination >5%.

***Fig. S6***.

Alluvial illustration of the fate of genomes in GMGC based on filters by GUNC and CheckM. Three filters are: 1) CheckM contamination <10%; 2) CheckM completeness <50%; 3) GUNC CSS <0.45 or GUNC contamination <2% (ignoring species level scores). The illustration shows the orthogonality and complementarity between GUNC and CheckM filters.

***Fig. S7***.

Mean F-score of 10 iterations of 10,000 non-chimeric vs 10,000 chimeric genomes across different values of GUNC CSS cutoffs used to separate between chimeric and non-chimeric genomes. For “All types” genome types 1 and 2 are used as non-chimeric and types 3, 4 and 5a are used as chimeric (type 5b excluded since it is not expected to be detected at all). For “No OOR” genome type 1 only is used as non-chimeric and types 3 and 4 are used as chimeric. The cutoff with high performance at “No OOR” was chosen so that its performance is as high as possible in the “All types” setup without any significant loss to “No OOR” setup performance.

***Fig. S8***.

Alluvial illustration of MAG “SRR1779121_bin.6” from Almeida et.al. 2019 that shows that GUNC can detect chimerism of related species when both source species have reference representation. The CheckM completeness is 79.31 and contamination is 1.72 for this MAG.

***Table S1***.

Contig lengths and coverages linked to gene-level taxonomy data underlying the visualization of genome in Fig. 3b.

