## Supplementary figures and images for "GUNC: Detection of Chimerism and Contamination in Prokaryotic Genomes"

### FigS1

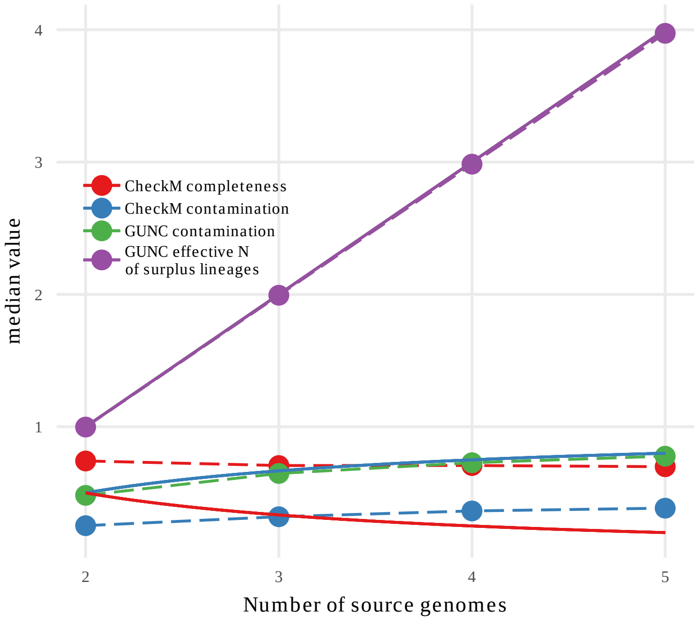

### FigS2

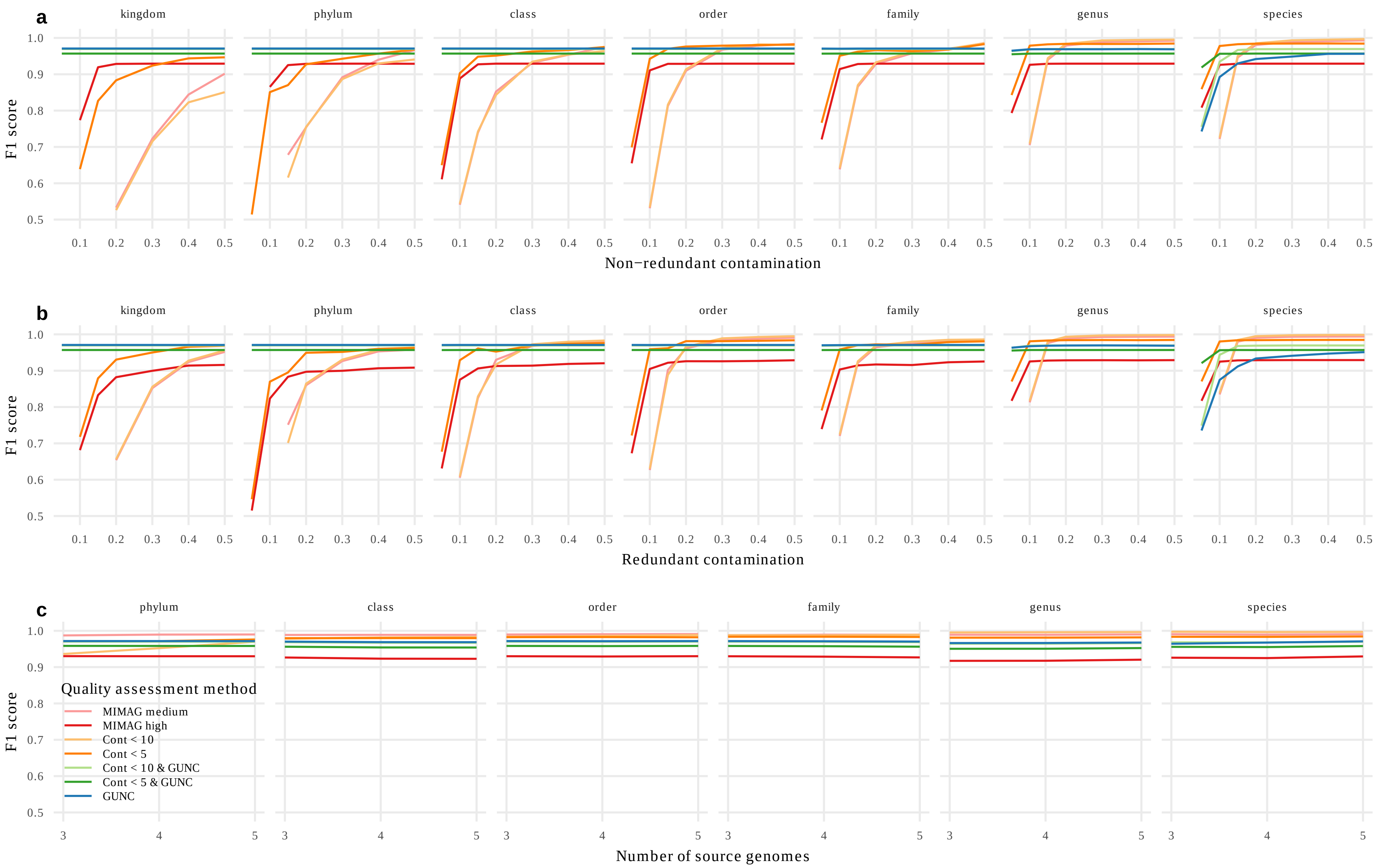

### FigS3

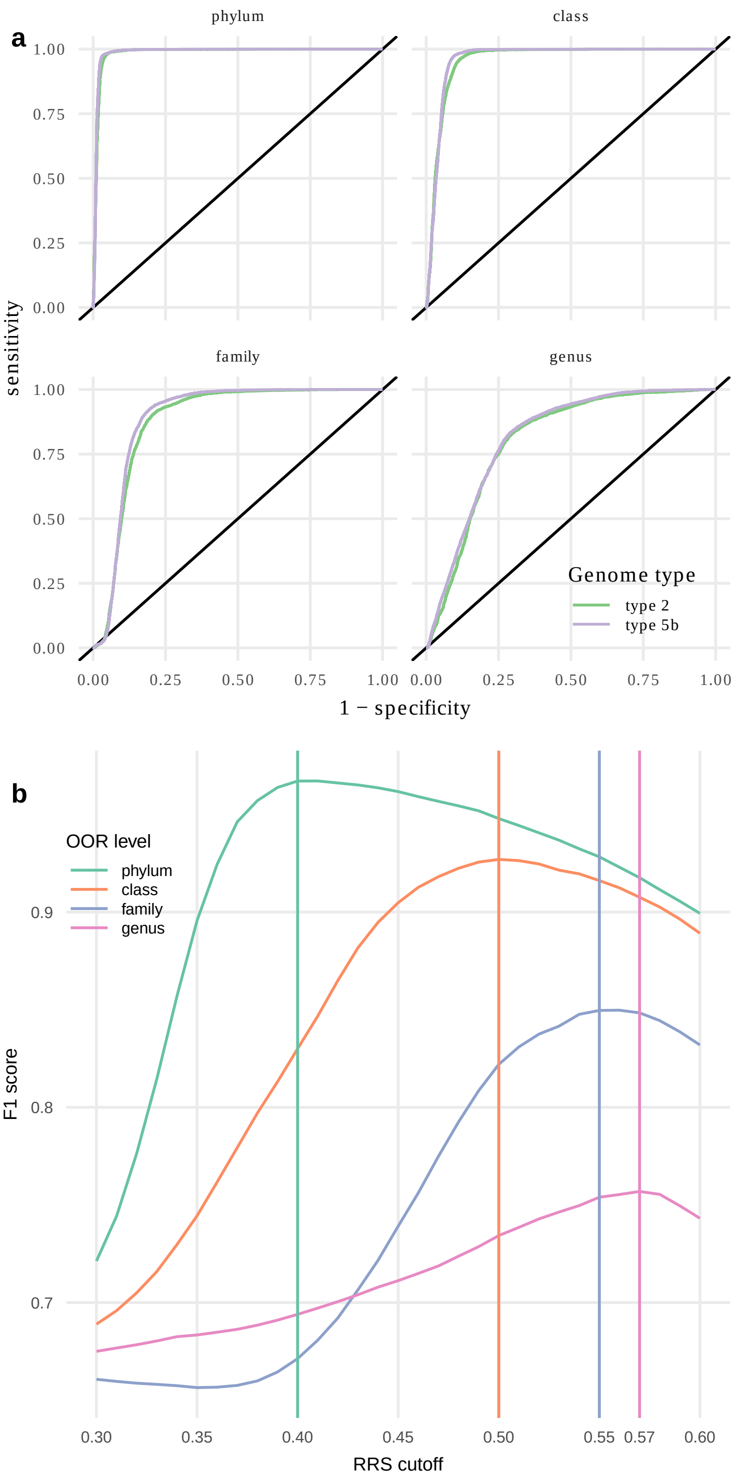

### FigS5

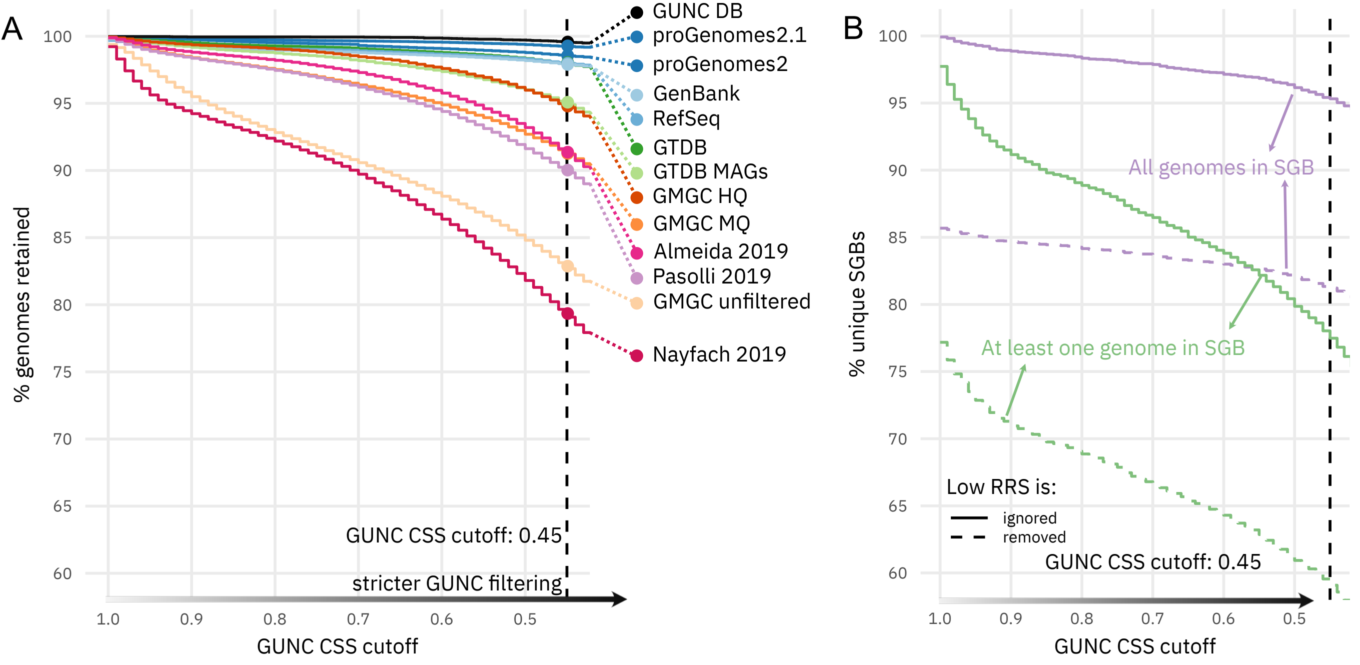

### FigS6

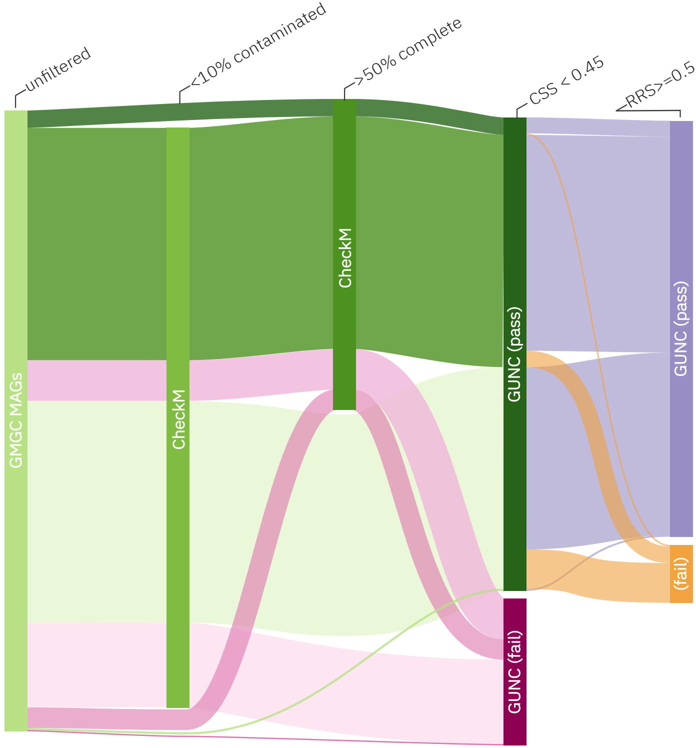

### FigS7

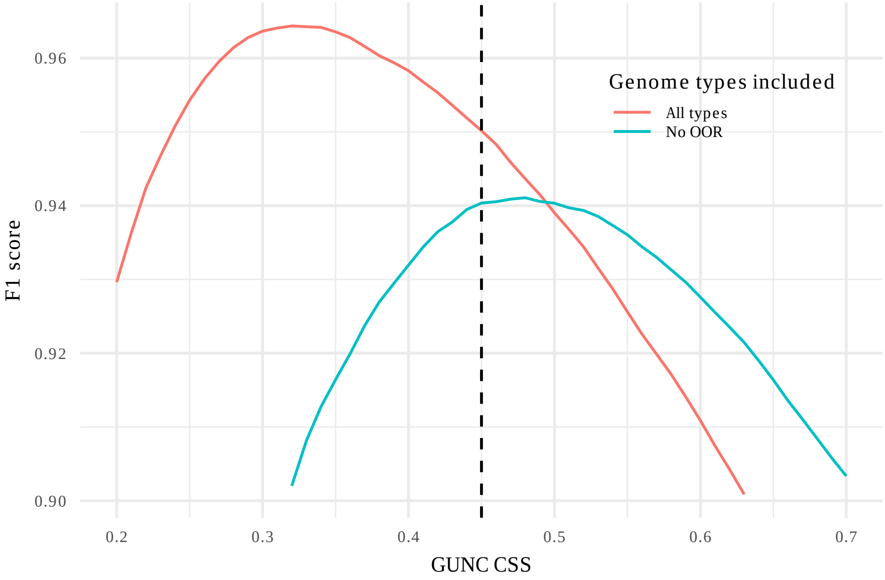

### FigS8

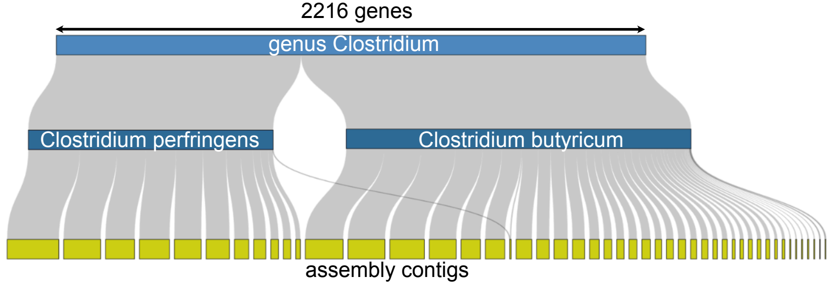
